# Optical Sectioning of Live Mammal with Near-Infrared Light Sheet

**DOI:** 10.1101/447433

**Authors:** Feifei Wang, Hao Wan, Jingying Yue, Mingxi Zhang, Zhuoran Ma, Qinchao Sun, Liangqiong Qu, Huilong Ma, Yeteng Zhong, Ye Tian, Guosong Hong, Wen Jung Li, Yongye Liang, Lianqing Liu, Hongjie Dai

## Abstract

Deep-tissue three-dimensional optical imaging of live mammals *in vivo* with high spatiotemporal resolution in non-invasive manners has been challenging due to light scattering. Here, we developed near-infrared (NIR) light sheet microscopy (LSM) with optical excitation and emission wavelengths up to ~ 1320 nm and ~ 1700 nm respectively, far into the NIR-II (1000-1700 nm) region for 3D optical sectioning through live tissues. Suppressed scattering of both excitation and emission photons allowed one-photon optical sectioning at ~ 2 mm depth in highly scattering brain tissues. NIR-II LSM enabled non-invasive *in vivo* imaging of live mice, revealing never-before-seen dynamic processes such as highly abnormal tumor microcirculation, and 3D molecular imaging of an important immune checkpoint protein, programmed-death ligand 1 (PD-L1) receptors at the single cell scale in tumors. *In vivo* two-color near-infrared light sheet sectioning enabled simultaneous volumetric imaging of tumor vasculatures and PD-L1 proteins in live mammals.

---

Optical imaging of biological systems capable of high spatiotemporal resolution *in vivo* and *ex vivo* has revolutionized biology and medicine for visualization and understanding of structures, functions and dynamic processes at the cellular and even molecular scale ^1,2^. To circumvent light scattering by tissues, *in vivo* 3D imaging by nonlinear two-photon fluorescence microscopy (670-1070 nm excitation) ^3-5^ or three-photon microscopy (1300-1700 nm excitation) ^6-9^ has reached penetration depths ~ 0.7-1.5 mm, benefiting from increased scattering mean free path of the near-infrared (NIR) excitation employed ^6^. Light sheet microscopy (LSM) uses orthogonally arranged planar illumination and wide-field detection, capable of high speed 3D optical sectioning, low photo-damage ^10,11^ and volumetrically imaging/tracking with subcellular resolution ^2^. Currently the excitation and emission of LSM are mostly in the visible except for two photon excitation in the NIR at ~ 940 nm ^12^ or three photon excitation at 1000 nm ^13^. Light scattering has limited LSM to imaging small transparent animals, organisms (zebrafish larvae, drosophila larvae, Medaka embryo, *C. elegans, etc.*), mammalian tissue samples after chemical clearing ^11,14,15^, and mouse brain at a depth of ~ 200 μm after craniotomy ^16^.

In recent years several classes of fluorescence probes have been developed with emission in the NIR-II window (1000-1700 nm) including carbon nanotubes, quantum dots, organic conjugated polymers and molecular dyes, and rare-earth nanoparticles ^17-30^. With suppressed photon scattering and diminished autofluorescence in the long-wavelength region, these probes have facilitated one-photon wide-field ^17-27^ or confocal ^28,29^ fluorescence imaging in the NIR-II window for mouse models of cardiovascular and brain diseases and cancer ^20,28,30^. Non-invasive imaging through the skin, skull and body tissues was achieved, with deep penetration depths and high signal-to-background ratio (SBR). Here, we developed NIR-II LSM using organic dyes and PbS/CdS core/shell quantum dot (CSQD) probes, extending excitation and emission to the unprecedented ~ 785-1320 nm and ~ 1000-1700 nm regimes respectively. Suppressed light scattering of both excitation and emission allowed up to 10 mm^3^ volumetric imaging of highly scattering mouse brain *ex vivo* with a penetration depth of ~ 2 mm. Importantly, NIR LSM readily afforded *in vivo* imaging of mouse tumor models non-invasively, enabling real-time observation of unusual tumor microcirculation, and 3D molecular imaging of immune checkpoint proteins at cellular scale in live mammals.

Our home-built LSM employed multiple switchable lasers with Gaussian beams (658 nm, 785 nm and 1319 nm) cylindrically focused into static light sheets for optical sectioning and an InGaAs camera for orthogonally detecting 900-1700 nm fluorescence (Fig. 1a and see setup details in Supplementary Fig. 1). We adjusted the effective numerical apertures of illumination objectives to produce light sheets with balanced waist thickness (~ 10-20 μm) and Rayleigh length (~ 0.5-2.0 mm) (see Methods for light sheet shape analysis), suitable for large scale volumetric imaging with single cell resolution. We employed several biocompatible NIR-II probes, an organic nanofluorophore p-FE ^28^ (excitation/emission: 650-850 nm/1000-1300 nm, Fig. 1b and Supplementary Fig. 2, dynamic light scattering size ~ 12 nm) and PEGylated PbS/CdS CSQD probes 29 (excitation/emission: ultraviolet to 1500 nm/1500-1700 nm, Fig. 1b and Supplementary Fig. 2, size ~ 6.9 nm). The two probes were sequentially injected intravenously into a mouse through the tail-vein at an interval of 5 min. We sacrificed the mouse at 30 min post injection while the probes were still circulating in the vasculature, fixed the brain and placed it in glycerol for *ex vivo* LSM imaging (see Methods for details). Under the same 785 nm light sheet (LS) excitation, we were able to clearly image the cerebral vasculatures as a function of tissue depth *Z* in three fluorescence emission windows 850-1000 nm (NIR-I, p-FE emission), 1100-1200 nm (NIR-IIa, p-FE emission) and 1500-1700 nm (NIR-IIb, CSQD emission) respectively (Fig. 1c and Supplementary Video 1). This allowed side-by-side comparison (Fig. 1c) of fluorescence LSM imaging in three sub-regions of 850-1700 nm under the same 785 nm LS excitation. Note that refractive index mismatch during scanning was compensated by linearly moving detection objective (Supplementary Figs 4 and 5) and no photobleaching was observed through this work, owing to the highly photo-stable NIR-II probes and minimal photo damage inherent to LSM ^10,11^.

**Figure 1.**
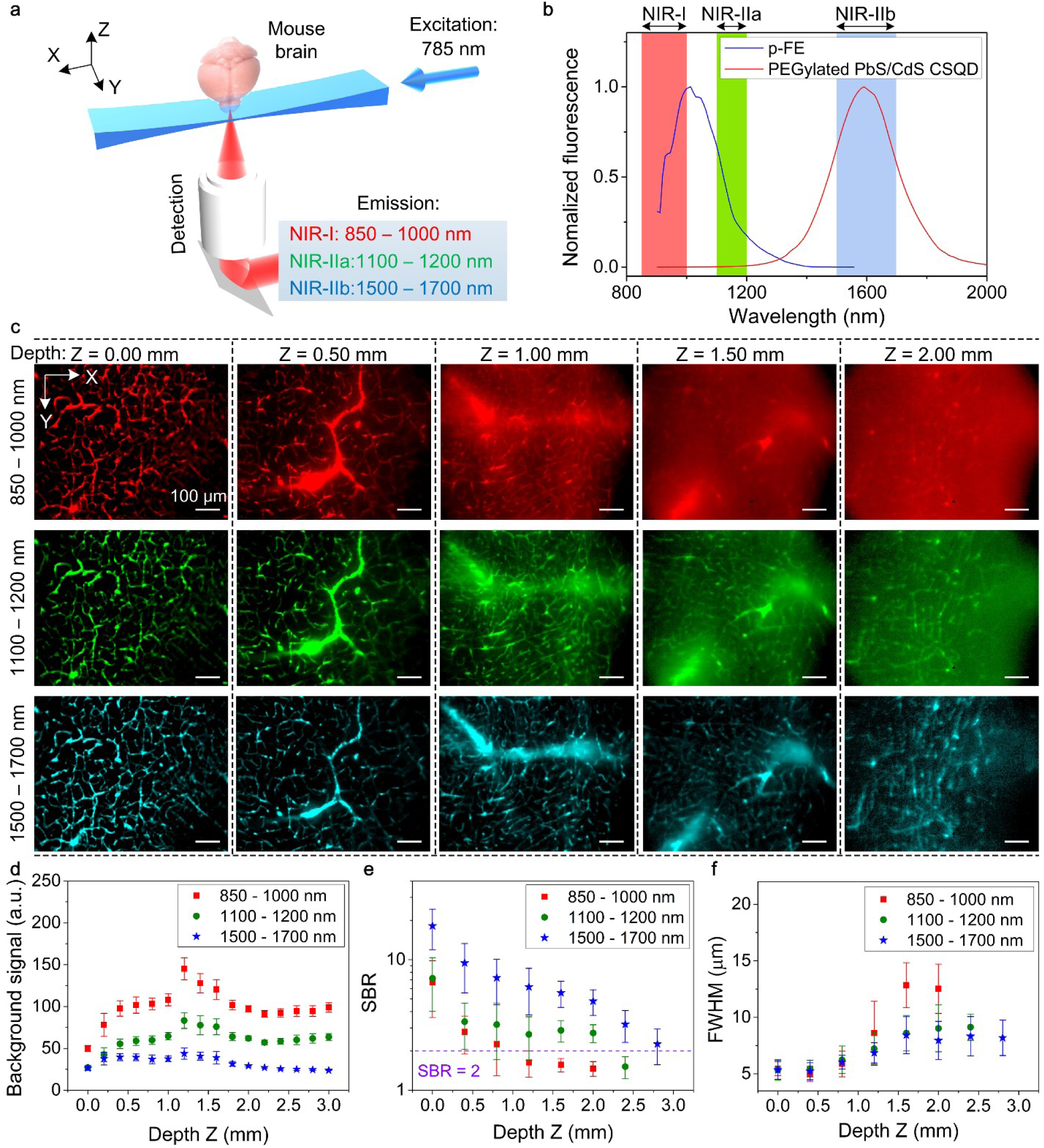
Light sheet microscopy in various NIR 850-1700 nm emission sub-regions. (**a**) A simplified schematic of the NIR LSM. (**b**) Fluorescence emission spectra of p-FE and PEGylated PbS/CdS core/shell quantum dots (see Supplementary Fig. 2 for excitation spectra). (**c**) Light-sheet optical sectioning mouse brain vasculatures at various depths in NIR-I, NIR-IIa and NIR-IIb emission regions using the same 785 nm light sheet illumination (also see Supplementary Video 1). The mouse brain tissue sample was prepared by intravenous injection of p-FE (emission: 850-1000 nm and 1100-1200 nm) and PEGylated PbS/CdS CSQD (emission: 1500-1700 nm) at 5-min interval. The mouse was scarified 30 min post injection of the probes while still in circulation. The mouse brain was taken out, fixed and preserved in glycerol for *ex vivo* imaging. See Supplementary Figs 4 and 5 for imaging details. The color bar range for each image is shown in Supplementary Fig. 3. Comparison of (**d**) background signal, (**e**) signal-to-background ratio and (**f**) FWHM of smallest vessels at various depths. Background was measured from randomly selected area without vasculatures. SBR is the ratio of fluorescence signals in randomly selected vasculatures over the background. (**d**-**f**) Error bars representing standard deviation were derived from analyzing ~ 10 target data at every depth. A 10X (NA = 0.25) imaging objective and a 5X illumination objective (effective NA = 0.039, light sheet waist *w* = ~15.2 μm and Rayleigh length *b* = ~ 1258.2 μm for 785-nm excitation, see Methods for light sheet shape analysis) were used in these experiments. Scale bars are 100 μm for all images in (**c**).

With a 785 nm light sheet, we observed that the brain tissue imaging depth limit increased (Fig. 1c and Supplementary Video 1), background signal decreased (Fig. 1d) and SBR increased (Fig. 1e) at longer detection wavelength from 850-1000 nm to 1100-1200 nm and 1500-1700 nm (Supplementary Video 1). The imaging depth limit (defined as tissue depth at which SBR decreased to ~ 2) increased from *Z*_SBR_=2 ~ 1.0 mm to ~ 2.0 mm and ~ 2.5 mm as emission wavelength increased from ~ 850 nm to ~ 1100 nm and ~ 1700 nm (Fig. 1e and Supplementary Video 1). These were direct results of suppressed scattering of emission photons (scattering ∝ *λ^-k^*, where *λ* is wavelength and *k* is in the range of 0.2–4.0 for biological tissues ^20^) under the same 785 nm excitation. Background signal caused by scattering was the lowest in the 1500-1700 nm emission range at all imaging depths (*Z* up to 3 mm, Fig. 1d and Supplementary Video 1). For emission in the 850-1200 nm range, background signal increased with tissue imaging depth up to *Z* = ~ 1 mm due to increased scattering by thicker tissue and decreased upon further increases in depth for *Z* > 1 mm (Fig. 1d). The latter was attributed to increased light absorption by thicker tissue that attenuated the background signal.

The lateral full width at half maximum (FWHM) values of the smallest cerebral vasculatures imaged in the three emission regions (850-1000 nm, 1100-1200 nm and 1500-1700 nm, Fig. 1f) at their tissue imaging depth limits of *Z*_SBR=2_ = 1.0 mm, 2.0 mm and 2.5 mm were ~ 7.2 μm, 9.0 μm and 8.3 μm respectively. Using an imaging objective with high magnification and numerical aperture (NA), < 5.0 μm lateral FWHM was achieved by NIR-LSM imaging by detecting 1500-1700 nm emission under a 785 nm LS illumination (Supplementary Fig. 6).

As a light sheet propagated in a scatting medium such as an intralipid phantom ^31,32^ mimicking the brain tissue, Monte Carlo simulations ^33^ and experiments observed light sheet decaying in intensity in the *X-Y* plane and spreading in *Z* due to tissue scattering (Fig. 2a,c,d, Supplementary Figs 7-10), which could hinder optical sectioning capability with reduced imaging field of view in *X-Y* and lower spatial resolution in *Z*. To circumvent this and maximize the benefit of reduced photon scattering at long wavelengths, we constructed a 1320 nm light sheet to afford the lowest degree of intensity decay and the least LS thickness broadening relative to the 785 nm and 658 nm light sheets (Fig. 2a,c,d, Supplementary Figs 8-10). In the fixed brain tissue, 658 nm, 785 nm and 1319 nm light sheets propagated over a distance of ~ 1.3 mm, ~ 1.7 mm and ~ 4.0 mm respectively (Fig. 2a), within which imaging of cerebral vasculatures by detecting 1500-1700 nm fluorescence of PbS/CdS core-shell quantum dots could still resolve small vessels (FWHM < 10 μm).

**Figure 2.**
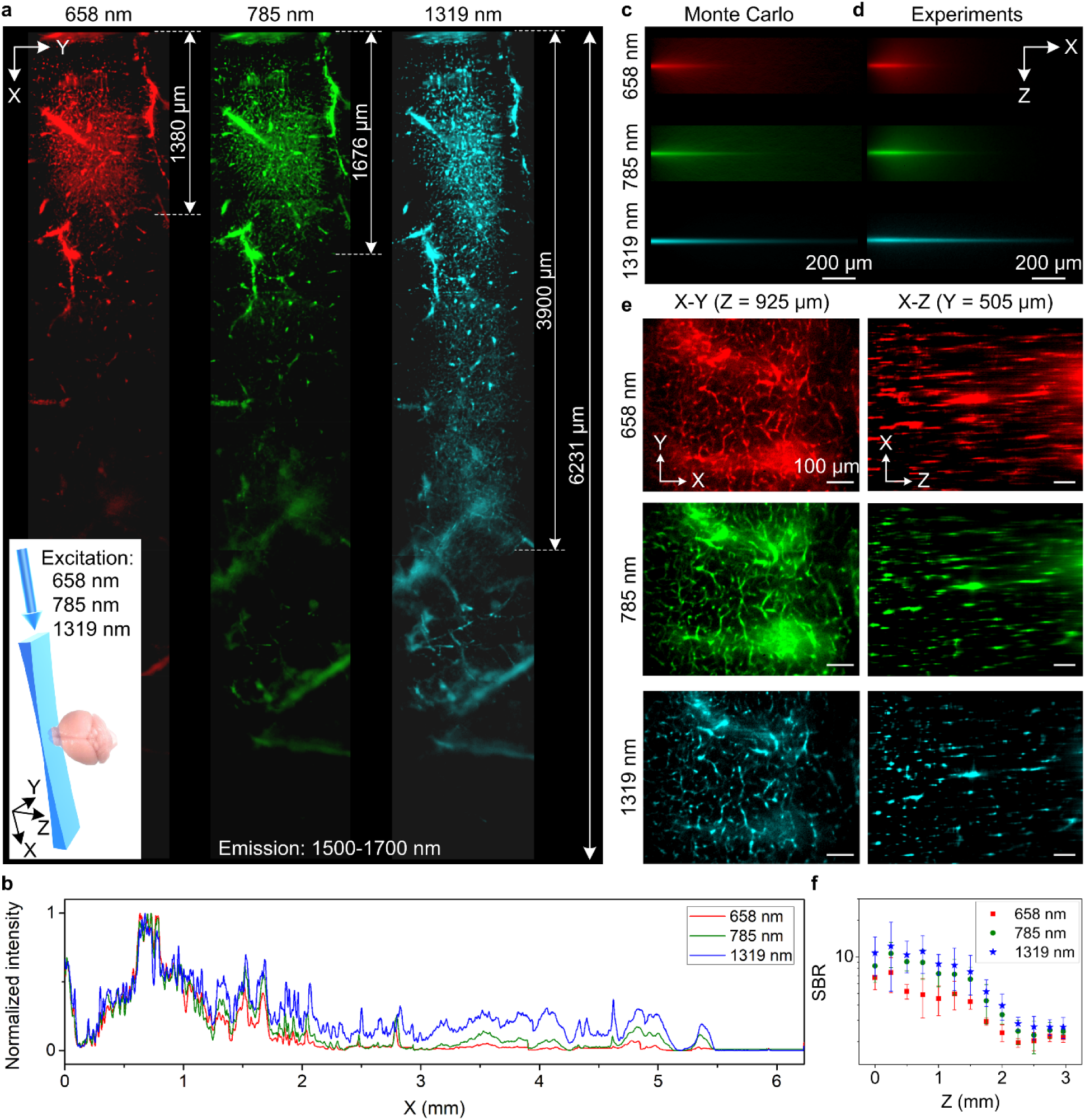
Propagation of light sheet excitation with progressively longer wavelength up to 1319 nm in brain tissues. (**a**) *X-Y* images of 1500-1700 nm quantum dot fluorescence in the vasculatures of a fixed brain tissue at a depth *Z* = ~ 200 μm under 658 nm, 785 nm and 1319 nm light sheet illumination as shown in the inset. The brain was taken out from a mouse 30 min after intravenous injection of PEGylated PbS/CdS CSQD, fixed and preserved in glycerol before imaging. 6 images were taken along *X* and stitched together for each light sheet. (**b**) Normalized sum intensity along *Y* direction of images in (**a**) as a function of propagation distance (*X*). (**c**) Monte Carlo simulations and (**d**) experimental results showing the *X-Z* propagations of different wavelengths light sheets in 2.5% intralipid tissue phantom (mimicking the brain) containing PEGylated PbS/CdS CSQD (emission: 1500-1700 nm). Scattering coefficient *μ*s = 109.3 cm^-1^, 73.5 cm^-1^ and 20.5 cm^-1^ and anisotropy *g* = 0.72, 0.64 and 0.34 were used to simulate 658 nm, 785 nm and 1319 nm excitation conditions respectively (see Supplementary Table 1). (**e**) Left: *X*-*Y* images of quantum dot 1500-1700 nm fluorescence in brain vasculatures taken at *Z* = 925 m under excitations by 658 nm, 785 nm and 1319 nm light sheets respectively (also see Supplementary Video 2). Right: images along the *X-Z* plane at a fixed *Y*, reconstructed from *X-Y* images at various depth *Z* (also see Supplementary Video 3). A 10X, 0.25-NA detection objective was used and LS excitation was generated by a 5X illumination objective with an effective NA of ~ 0.039 (see Methods for experimental details and Supplementary Fig. 7 for light sheet shape analysis). (**f**) Comparison of SBR for *X*-*Y* images recorded at different depth for 658 nm, 785 nm and 1319 nm excitation. About 10 randomly selected vasculatures and 10 areas without vasculatures were analyzed to calculate SBR at each depth. Error bars represent standard deviation. Scale bars, 200 μm (**c**,**d**) and 100 μm (**e**).

It constituted a breakthrough in one-photon imaging by exploiting long wavelength excitations up to 1300 nm (Supplementary Fig. 2). Remarkably, the 1319 nm light sheet could propagate > 6 mm to allow imaging of large blood vessels in the highly scattering mouse brain over large field of views (Fig. 2a). Suppressed scattering of longer wavelength LS was also gleaned from *X-Y*, *X-Z* or *Y-Z* cross sectional images (Fig. 2e, Supplementary Fig. 11 and Supplementary Videos 2 and 3), with improved SBR (Fig. 2f) and reduced FWHM of feature sizes along the depth *Z* direction, corresponding to higher vertical resolution and better sectioning capability along *Z* (Supplementary Fig. 11e,f).

Light sheet microscopy with both excitation and emission in the 1000-1700 nm NIR-II window minimized scattering and maximized the penetration depth and field of view. High resolution 3D NIR-II LSM sectioning (Fig. 3a and Supplementary Video 4, volume = 810 μm x 648 μm x 3000 μm, 3 μm *Z* increment in depth) afforded sub-10 μm x 10 μm x 15 μm volumetric resolution (FWHM) (Fig. 1f and Supplementary Fig. 11e,f).

**Figure 3.**
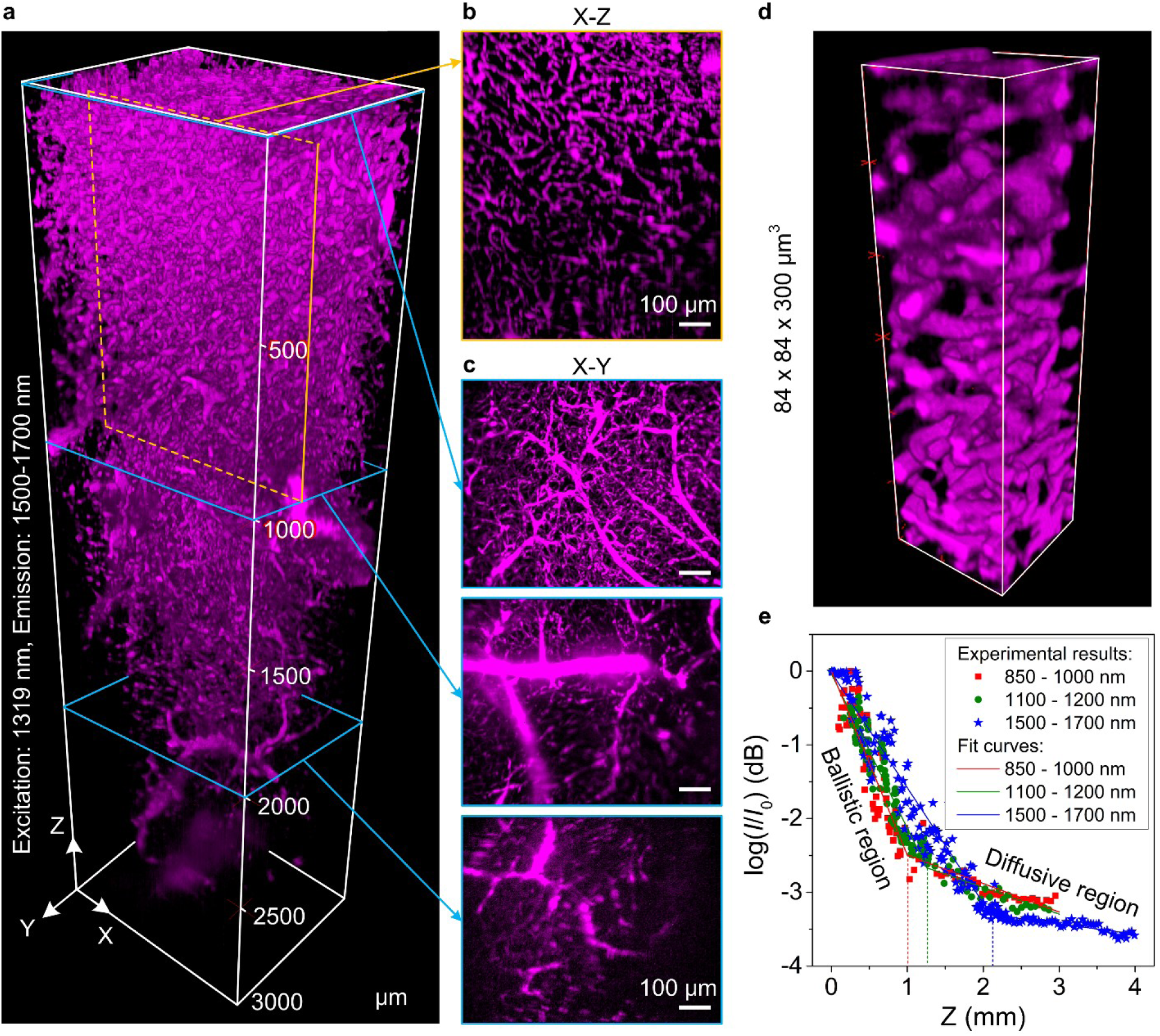
Volumetric 1500-1700 nm fluorescence imaging of mouse brain sectioned by a 1319 nm light sheet. (**a**) 3D rendering of PEGylated PbS/CdS CSQD labelled vasculatures in mouse brain. 1500-1700 nm fluorescence was collected at 1319 nm excitation by a 10x detection objective. A 5x illumination objective (with an effective NA = ~ 0.051) was used to generate LS excitation (also see Supplementary Video 4). The scanning increment along *Z* was 3 μm. The excitation power (~ 8 mW) and the exposure time (0.8 s) were kept constant during entire sectioning. (**b**) Maximum-intensity *Y*-projection (50 μm in thickness along *Y*, the maximum-intensity *Y*-projection took the brightest pixel in *X-Z* layers through 50-μm *Y* distance and displayed the maximum intensity values in the final 2D *X-Z* image) and (**c**) maximum-intensity *Z*-projections for a 150 μm-thick volume along *Z* at *Z* = 0 μm, 1000 μm and 2000 μm, respectively. (**d)** 3D rendering of a smaller region in (**a**). (**e**) Maximum fluorescence intensity (*I*) in different emission regions detected at various depths (*Z*) in the mouse brain. *I*_0_ is the fluorescence intensity at *Z* = 0 μm. In these experiments, the brain was taken out from a mouse intravenously injected with p-FE (excitation: 785 nm, emission: 850-1000 nm and 1100-1200 nm) and PEGylated PbS/CdS CSQD (excitation:785, emission: 1500-1700 nm) with 5-min interval at 30 min post injection. Then the mouse brain was fixed and preserved in glycerol for *ex vivo* imaging. Scale bars, 100 μm (**b**,**c**).

Under the 785 nm LS excitation, the maximum 1500-1700 nm fluorescence signal detected in the cerebral vasculatures of mouse brain cortex layer as a function of depth *Z* showed two attenuation regions (Fig. 3e). There was an initial exponential attenuation due to reduction in ballistic and nearly ballistic photons (slightly deflected) emerging through the tissue following the Beer–Lambert law *I*_ball_ = *I*_0_ e^-*z*/*ls*^ (where *z* is imaging depth, *I*_0_ is initial intensity at *z* = 0 mm and *l_s_* is scattering mean free path MFP). This was followed by a slower decay region at deeper *Z* from which multiply scattered photons diffusing through the brain tissue (diffusive region) were dominant ^34^. The MFP *l_s_* extracted (Fig. 3e) was ~ 400 μm, ~ 477 μm and ~ 639 μm for 850-1000 nm, 1100-1200 nm, 1500-1700 nm windows respectively. The imaging penetration depth limits (see Supplementary Table 1 for detailed scattering coefficients and MFP comparison) were ~ 2.5*l_s_* (for 850-1000 nm emission), ~ 2.6*l_s_* (for 1100-1200 nm emission) and ~ 3.3*l_s_* (for 1500-1700 nm emission).

The capability of NIR-II LSM performing volumetric imaging through scattering tissues at the 1-10 mm^3^ scale enabled non-invasive *in vivo* 3D imaging of protruding features on live mice related to disease models (Fig. 4b), facilitating cellular resolution LS sectioning through intact tissues for mammals. We carried out NIR LSM imaging of subcutaneous xenograft tumors on mice ear and right/left flank of back without invasive surgery or installing optical windows ^35^. *In vivo* hemodynamic imaging of a tumor model used for immunotherapy, murine colorectal MC38 tumors on C57BL/6 mice ear was carried out with the NIR-II light sheet fixed at a *Z* position ~ 300 μm below the top (where the fluorescence signal was first detected) of the tumor (~ 8 mm in diameter, Fig. 4a,b). Time-course LS imaging of the p-FE nanofluorophore immediately following intravenous injection into the mouse tail-vein (785 nm excitation, 1000 nm detection at exposure times ~ 100-800 ms) revealed abnormal microcirculation in tumor. Blood flows in tumor vasculatures were found irregular and intermittent (Fig. 4a,c and Supplementary Video 5), with turning-on and shutting-off behavior, oscillatory/fluctuating flowing patterns and even flow direction reversal in the same vasculature (Fig. 4c, marked by arrows, Supplementary Video 5).

**Figure 4.**
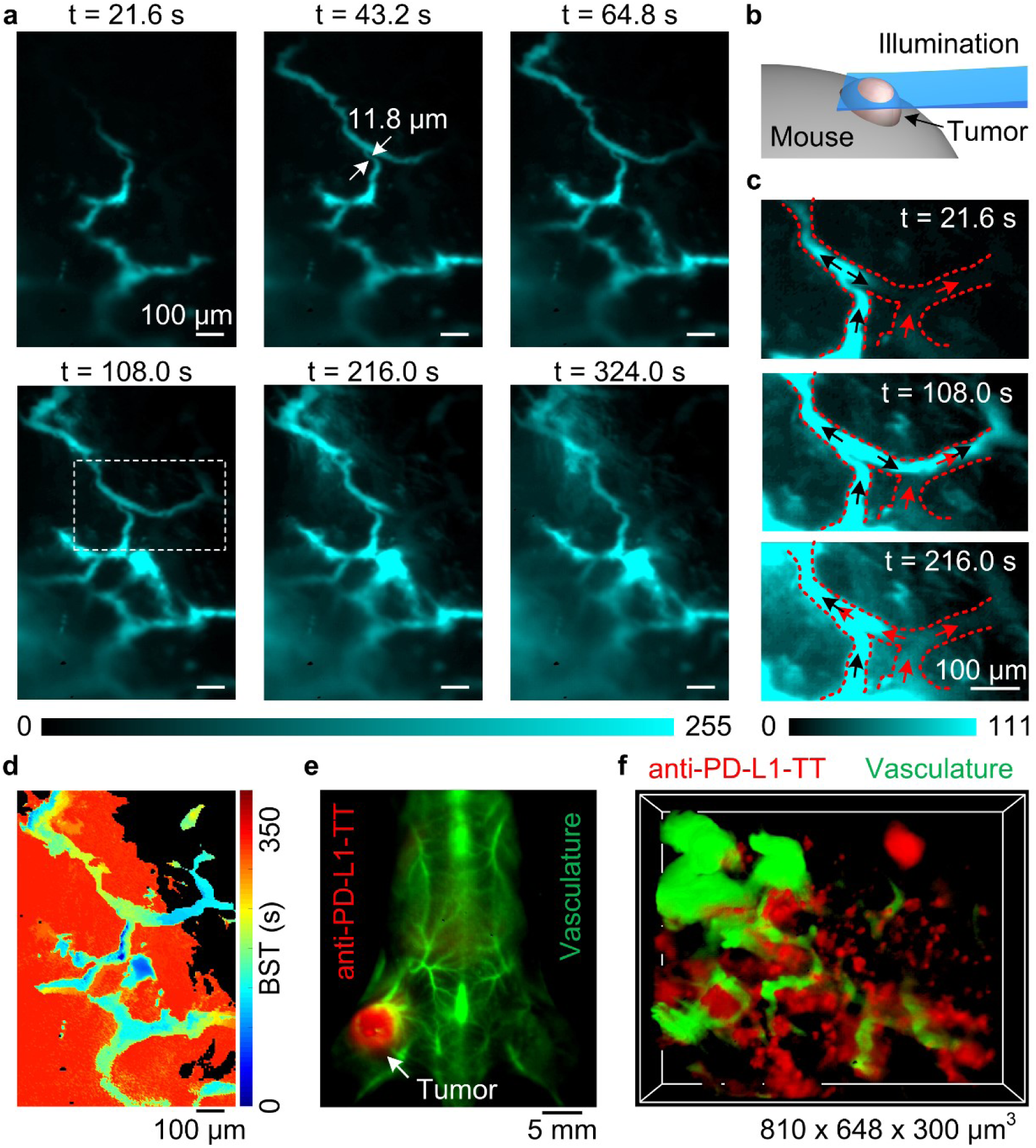
*In vivo* NIR-II light sheet microscopy of xenograft tumors in mice. (**a**) Time-course (Supplementary Video 5) LSM of tumor vasculatures at a fixed illumination plane below the top of a xenograft MC38 tumor at *Z* ~ 300 μm after intravenous injection of p-FE (excitation: 785 nm, emission: 1000-1200 nm). A 4X detection objective and a 5X illumination objective were used. (**b**) Schematic illustration of *in vivo* NIR LSM. (**c**) Abnormal blood flows in tumor vessels, showing on-off intermittency and direction reversal in the rectangular highlighted region in (**A**) and gradual extravasation into tumor space (Supplementary Video 5). Black arrows and red arrows represent blood with or without p-FE respectively. (**d**) A BST map (blood supply time, defined as the time at which the pixel intensity reached its maximum value relative to the fluorescence flow is first detected) showing highly heterogeneous blood perfusion in tumor vessels and slow, inhomogeneous extravasation behavior into tumor space. (**e**) Wide-field imaging of a mouse. The anti-PD-L1-TT probes (red color, excitation: 808 nm, emission: 1000-1200 nm, exposure time 5 ms) were injected intravenously and kept circulation for 24 h. 30 min prior to tumor imaging PEGylated PbS/CdS CSQD (green color, excitation: 808 nm, emission: 1500-1700 nm, exposure: 5 ms) was injected and imaged for overlaying with the tumor specifically labeled by the anti-PD-L1-TT dye. (**f)** *In vivo* two-color 3D light sheet microscopy of anti-PD-L1-TT (red color, excitation: 785 nm, emission: 1000-1200 nm, exposure: 0.8 s) and vasculatures (green color, PEGylated PbS/CdS CSQD, excitation: 1319 nm, emission: 1500-1700 nm, exposure: 0.8 s) in a MC38 tumor using a 10X detection objective and a 5X illumination objective (Supplementary Video 6). The *Z* scanning increment was 3 μm. The discrete red spots were down to 6 x 6 x 15 μm^3^ in size, corresponding to PD-L1 expressing cells inside the tumor. No such spots were observed in tumor injected with TT dye without any anti-PD-L1 conjugated (Supplementary Fig. 13a). Scale bars, 100 μm (**a**, **c**, **d**); 5 mm (**e**).

Intermittent blood flows was only inferred from *ex vivo* histological methods previously ^36^, suggesting unstable pressure within a tumor due to uncontrolled tumor angiogenesis. Further, blood supply time (BST) analysis revealed that the p-FE nanofluorophore gradually leaked out from some vasculatures (in the red-colored region in Fig. 4d) but not from others (in the black-colored region in Fig. 4d).

To further exploit NIR-II LSM, we performed *in vivo* two-color molecular imaging and vasculature imaging of PD-L1 expressing MC38 tumor ^37^ using a renal excretable organic dye ^38,39^ (TT dye: excitation ~785 nm/emission ~1000-1200 nm) conjugated to anti-PD-L1 antibodies and unconjugated PEGylated PbS/CdS CSQD (excitation ~ 1319 nm/emission 1500-1700 nm) intravenously injected into tumor bearing mice. PD-L1 is an important immune checkpoint protein expressed by tumors as a powerful way of T cell immunity evasion. PD-L1 blocking by antibody immunotherapy is a novel approach to treat various cancers. *In vivo* PD-L1 imaging is important to fundamental understanding of cancer immunity, and to immunotherapy prognosis since treatment efficacy correlated with PD-L1 levels in the tumor ^40,41^. 24 h post injection of anti-PD-L1-TT dye, we first performed wide-field imaging and observed much brighter anti-PD-L1-TT dye signals in MC38 tumors ^37^ (Fig. 4e) than post injection of the unconjugated free TT dye (Supplementary Fig. 12b). Since wide-field imaging only provided 2D projected signals and lacked spatial resolution, we switched to 3D *in vivo* NIR LSM to profile PD-L1 receptors at various depths of the tumor (Fig. 4f) inside the live mouse, a task only done with tumor biopsies previously ^42-44^. We observed discrete anti-PD-L1-TT dye labelled features inside tumors with sub-6-μm FWHM in the lateral *X-Y* plane and sub-15-μm FWHM in *Z* using a 50X detection objective (Supplementary Fig. 13b,d), suggesting cellular scale molecular imaging of PD-L1 *in vivo*. Meanwhile, we imaged irregular tumor blood vessels engulfing PD-L1 expressing cells upon intravenously injecting PEGylated PbS/CdS CSQD (excitation 1319 nm/emission 1500-1700 nm) circulating in the vasculatures with sub-5 μm x 5 μm x 10 μm volumetric resolution (FWHM) (Supplementary Fig. 13c,e). The results suggested tumor extravasation of anti-PD-L1-TT dye (injected 24 h prior to CSQD injection) and specific targeting PD-L1 expressing cells in the tumor of live mouse.

This work developed 3D near-infrared light sheet microscopy for *in vivo* and *ex vivo* deep tissue volumetric imaging through highly scattering biological tissues. Light sheet microscopy with both excitation and emission in the NIR-II window avoided shadows or stripes along the illumination direction by suppressing tissue scattering and adsorption effects encountered by visible LSM ^45^. Non-invasive NIR-II LSM enabled *in vivo* observation in wide-field detection mode with suppressed background, which facilitates dynamic processes tracking and molecular imaging at cellular resolution over millimeter scale simultaneously and could provide a complementary method to two-photon microscopy with lower cost and under less invasive conditions. NIR LSM imaging can be further advanced by developing brighter fluorophores with added colors and applying new configurations of LSM ^46^. Recent developments such as noncoherent structured illumination ^47^ and optical lattices illumination ^2^ could be introduced to improve the resolution and contrast of NIR-II LSM. Real time molecular imaging of multiple targets by rapid sectioning 3D tissues of live mammals should become possible.

## Methods

### NIR-II fluorescence probes

This work used an organic nanofluorophore p-FE dye, PEGylated core-shell quantum dots PbS/CdS CSQD (see Supplementary Fig. 2), and an organic renal excretable TT dye. p-FE is comprised of organic dyes trapped in amphiphilic polymeric micelles ~ 12 nm in size measured by dynamic light scattering ^28^. The PEGylated PbS/CdS CSQD was developed recently exhibiting a wide range of excitation wavelength spanning from UV to ~ 1300 nm, high brightness, biocompatibility and liver excretion ^29^. The TT dye ^38,39^ exhibits similar spectroscopic properties as the p-FE dye with an excitation ~ 785 nm and emission ~1000-1200 nm. The TT dye contains an azide group that can be used for conjugation to PD-L1 antibodies by click chemistry for molecular imaging ^38,39^. Purified anti-mouse CD274 (B7-H1, PD-L1) antibody was purchased from GoInVivo (Biolegend, Cat#124328). Conjugation was done through copper-free click chemistry using linker DBCO-PEG4-NHS purchased from Click Chemistry Tools ^48^.

### NIR-II light sheet microscope setup

As show in Supplementary Fig. 1, a laser (Gaussian beam) with wavelength of 658 nm, 785 nm or 1319 nm with maximum excitation power of 1.7 mW, 11.9 mW and 8.2 mW respectively was directed into a spatial filter consisting two achromatic lenses (L3 and L4) and a 50-μm pinhole (PH). This spatial filter was introduced to improve the circularity and quality of the illumination beam and to generate uniform light sheet across the field of view. For in vivo imaging, the actual excitation intensity illuminated on mouse was ~ 65 mW cm^-2^, ~ 174 mW cm^-2^and ~ 89 mW cm^-2^ for 658 nm, 785 nm and 1319 nm laser, which is below the safety limit for laser exposure (658 nm: 200 mW cm^-2^; 785 nm: 300 mW cm^-2^, 1319 nm: 1000 mW cm^-2^)49. The excitations could be selected by removable mirrors (M2-M4). Excitation power was measured by a laser power meter (3A-SH, NOVA II, OPHIR). Before the laser entering a cylindrical lens (CL), a vertically arranged adjustable mechanical slit parallel to the CL was used to adjust the span range of light sheet along *Y*-axis direction (Supplementary Fig. 1). A pair of achromatic lenses (L1, L2) and another slit (S1) was used for adjusting the effective numerical aperture of the objective (for adjusting light sheet shape, see below) before the light was focused on the back focal plane of illumination objective (O1) to form the light sheet illumination.

The light sheet was positioned to pass through a tissue sample or a tumor protrusion on a mouse, fluorescence imaging was done by focusing on and normal to the light sheet plane through a detection objective (O2) and a 200-mm tube lens, using a liquid-nitrogen-cooled InGaAs camera (2D-OMA V, Princeton Instruments) after filtered by selected emission filters. The focal lengths of L1, L2, L3, L4 were 60 mm, 100 mm, 30 mm and 60 mm, respectively. All the optical components were made by Thorlabs. For illumination objective, we used a 5X objective (NA = 0.15, Nikon LU Plan) or a 10X objective (NA = 0.25, Bausch & Lomb Optical Co.). For imaging, we used a 4X objective (NA = 0.1, Bausch & Lomb Optical Co.), a 10X objective (NA = 0.25, Bausch & Lomb Optical Co.) or a 50X objective (NA = 0.6, Nikon CF Plan) (see Supplementary Table 2 for combinations of objectives for various experiments).

### Light sheet shape, resolution and field of view considerations

Two orthogonal slits were mounted in the illumination arm to adjust the light sheet shape by changing the effective numerical aperture NA and the span range along *Y*-axis direction (Supplementary Fig. 1). To observe the light sheet propagation in glycerol, water, intralipid or brain, the cylindrical lens was rotated by 90o and S1 was adjusted to control the light sheet spanning along vertical direction and S2 was used to control the actual numerical aperture (NA) of illumination. This allowed us to form light sheets with adjustable waist thickness and Raleigh length to balance imaging resolution and field of view (FOV). When 10X illumination objective and 50X detection objective were used, the effective NA of illumination was adjusted to be ~ 0.17. For 5X illumination objective and 4X or 10X detection objective, the effective NA of illumination was adjusted to be ~ 0.039 or ~ 0.051.

The effective NA was estimated using NA = *n* sin*α* = *n* sin(arctan(*D*/2*f*)), where *n* is the refractive index, *α* is the half of aperture angle, *D* is the illumination width of light sheet (adjusted by slit S1 for LSM imaging) as it exiting the illumination objective, and *f* is the focal length of illumination objective. At a given width of slit S1, *D* was measured experimentally by putting a scattering paper close to the aperture of the illumination objective. *D* was adjusted by S2 when the cylindrical lens was rotated by 90o for imaging the side view of the light sheet (Supplementary Fig. 7). The measured light sheet waist and Rayleigh length were consistent with those from theoretical estimations based on the effective NA values (see Supplementary Fig. 7).

The diffraction limited resolution along *Z* for 10X, 0.25-NA detection objectives was 14.8 μm (850-1000 nm), 18.4 μm (1100-1200 nm) and 25.6 μm (1500-1700 nm) estimated by *nλ*/NA^2^ ^50^. The value (25.6 μm) in 1500-1700 nm window was larger than the measured LS thickness (Supplementary Fig. 7). Therefore, the resolution along *Z* was dominant by LS thickness under this condition ^10^, which is down to ~ 10 μm (Supplementary Fig. 7). Using the 0.5-NA 50X detection objective, the diffraction limited resolution along *Z* was 2.6 μm (850-1000 nm), 3.2 μm (1100-1200 nm), 4.4 μm (1500-1700 nm). These analysis suggested that the Z resolution of our current NIR-II LSM is down to ~ 10 μm, suitable for resolution along *Z* at the single cell level. *X*-*Y* diffraction limited resolution is higher, ~ 2.3 μm (850-1000 nm), 2.8 μm (1100-1200 nm) and 3.9 μm (1500-1700 nm) for the 10X, 0.25-NA detection objective and 0.9 μm (850-1000 nm), 1.2 μm (1100-1200 nm) and 1.6 μm (1500-1700 nm) for the 50X, 0.6-NA detection objective (estimated by Rayleigh criteria, 0.61*λ*/NA).

The LS waist and Rayleigh length are contradicting factors, optimizing one means degrading performance in the other. The selecting actual NA for each experiment were tradeoffs of these two parameters to obtain uniform LS that are as thin as possible across a large enough FOV. The actual NA and corresponding waist and confocal length for each data set were summarized in Supplementary Table 2.

### Light sheet microscopy 3D volumetric imaging/scanning

For 3D NIR-II LSM imaging, as the imaging depth changed, an obvious misalignment of light sheet and working plane of the imaging objective appeared due to refractive index mismatch of the air and tissue. This was compensated by a linear movement of detection objective (Supplementary Figs 4 and 5) concurrent with sample Z position change. The sample scanning in *Z* and detection objective *Z* compensation movement was realized by a 3D translation stage (KMTS50E, Thorlabs) and a single-axis translation stage (MTS50-Z8, Thorlabs), respectively (see Supplementary Fig. 4). Different detection windows were selected by using corresponding long-pass and short-pass filters. Synchronous control of 3D translation stage movement and image record was realized using LabView software through a data acquisition card (NI USB-6008, National Instruments). Maximum intensity projections (Fig. 3b,c) and 3D rendering was performed using ImageJ. Multi-color fluorescence images were also merged in ImageJ ^51^.

### Mouse handling and tumor xenograft

Mouse handling was approved by Stanford University’s administrative panel on Laboratory Animal Care. All experiments were performed according to the National Institutes of Health Guide for the Care and Use of Laboratory Animals. C57BL/6 female mice were purchased from Charles River. Bedding, nesting material, food and water were provided. 6-week-old C57BL/6 mice were shaved using hair removal lotion and inoculated with ~ 1 million MC38 cancer cells on the right flank of back near the hindlimb or on the ear for tumor growth. The sample sizes of mice were selected based on previously reported studies. Mice were randomly selected from cages for all experiments. All relevant data are available from authors. During *in vivo* imaging, all mice were anaesthetized by a rodent anesthesia machine with 2 l min^-1^ O2 gas mixed with 3 % isoflurane.

### *Ex vivo* NIR light sheet microscopy of mouse brains

For *ex vivo* LSM of mouse brains in various NIR-I and NIR-II regions (data in Fig. 1c, Fig. 2e, Supplementary Figs 4, 5, 6 and 11), C57BL/6 mice were firstly injected with 200 μL of p-FE with O.D. = 4 at 808 nm, followed by injection of 200 μL PEGylated PbS/CdS CSQD (O.D. = 4 at 808 nm) at 5 min post injection of p-FE. The mouse was sacrificed at 30 min post injection under anesthesia, and brain tissues were taken out and fixed with 10% neutral buffered formalin at room temperature. After washing in PBS twice, the fixed mouse brain was preserved in glycerol at 4 °C for further imaging.

For NIR-II imaging mouse brains (data in Fig. 2a and Fig. 3a), C57BL/6 mice were injected with 200 μL PEGylated PbS/CdS CSQD (O.D. = 4 at 808 nm) and sacrificed at 30 min post injection. The brain tissues were taken out and fixed with 10% neutral buffered formalin at room temperature. After washed in PBS twice, the fixed mouse brain was preserved in glycerol at 4 °C for *ex vivo* imaging.

For 5X illumination objective and 4X (for data in Fig. 2a) or 10X (for data in Fig. 1c, Fig. 2e, Supplementary Figs 4, 5 and 11) detection objective, the actual NA of illumination was adjusted to be ~ 0.039 or ~ 0.051. When 10X illumination objective and 50X detection objective were used, the actual NA of illumination was adjusted to be ~ 0.17 (Supplementary Fig. 6). The corresponding waists for different wavelengths were shown in Supplementary Fig. 7. Other detailed experimental conditions, such as *Z* scanning increment, exposure time, excitation and emission wavelengths, were summarized in Supplementary Table 2.

### *In vivo* wide-field NIR-II fluorescence imaging

All NIR-II wide-field fluorescence images (data in Fig. 4e and Supplementary Fig. 12) were recorded using a 2D liquid-nitrogen cooled InGaAs camera (Princeton Instruments, 2D OMA-V, USA). An 808 nm fiber-coupled diode laser (RPMC Lasers, USA) was used as the excitation source and a filter set (850 and 1,000 nm short-pass filter) was applied to filter the excitation light. The actual excitation intensity after passing filters was ~ 70 mW cm^-2^. The fluorescence signal was collected by two achromatic lenses to the InGaAs camera with different magnifications after filtered by corresponding low-pass and long-pass filters. Two-channel fluorescence images were merged in ImageJ.

### *In vivo* NIR light sheet microscopy of tumors

C57BL/6 mouse bearing a xenograft MC38 tumor on the ear or right/left flank of the back near the hindlimb was injected with anti-PD-L1-TT dye or free TT dye. The mouse was used for *in vivo* light sheet microscopy imaging immediately and at 24 hour post injection by placing the mouse on a 3D translation stage (KMTS50E, Thorlabs) after anesthesia. The initial LS illumination position *Z* below the top of the surface of the protruding tumor was controlled by the 3D motorized translation stage. For monitoring dynamic blood flow at a fixed illumination plane through the tumor (data in Fig. 4a), the camera began recording with a preset exposure time immediately after p-FE (200 μL, O.D. = 4 at 808 nm) or PEGylated PbS/CdS CSQD (200 μL, O.D. = 4 at 808 nm) was injected into the tail vein. When the recorded fluorescence images did not show further changes and reached a steady state, 3D light sheet microscopy was performed to volumetrically imaging the vasculatures or mapping the distribution of anti-PD-L1-TT in the tumor (for Fig. 4f). For dynamic observation of blood flow (Fig. 4a, Supplementary Video 5), a 4X imaging objective lens was used. The illumination was generated by a 5X illumination objective with actual NA of ~ 0.039 (Supplementary Fig. 7). The PD-L1 receptors were profiled using a 10X (Fig. 4f) and a 50X (Supplementary Fig. 13b-e) detection objective. The *Z* scanning increment, exposure time and excitation and emission wavelengths were summarized in Supplementary Table 2.

### Study of light sheet propagation in different media

We experimentally compared light sheet propagation in glycerol solutions using light sheets with different NA and excitation wavelengths. These experiments were performed in glycerol containing PEGylated PbS/CdS CSQD uniformly dispersed in glycerol. The emission was collected in 1500 – 1700-nm window excited by 658-nm, 785-nm and 1319-nm LS illuminations. In order to directly observe light transmission in glycerol, we rotated the cylindrical lens by 90° and used mechanical slits to control the actual NA and the spanning range along *Y* (Supplementary Fig. 1). By so doing, the illumination plane was rotated by 90° and the light sheet shape can be imaged along the *Y* direction for its X-Z plane for side view (Supplementary Fig. 7a). In a transparent medium (Supplementary Fig. 7a), our experimentally measured waist and double Rayleigh range of light sheet were consistent with theoretical estimations (Supplementary Fig. 7b,c).

To study the propagation of the light sheet with wavelength of 658 nm, 785 nm and 1319 nm in a scattering medium, we performed experimental imaging of LS propagation in intralipid solutions of different concentrations (Supplementary Fig. 8a). Light scattering was apparent as the intralipid concentration increased from 0.00% to 5.00 % when 658-nm or 785-nm excitation was used. Impressively, 1319-nm LS excitation retained its shape over the longest distance. We further simulated the light sheet propagation in the intralipid phantom by Monte Carlo method based on the method developed by Wang *et al*. ^33^ using the scattering coefficient *μ*s and the anisotropy *g* estimated by ^31,32^:

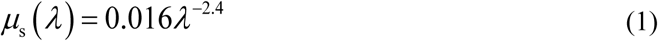

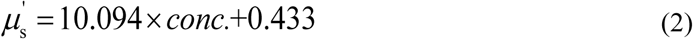

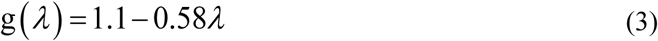

Where *λ* is the wavelength, *μ*s’ = *μ*_s_(1-*g*) is the reduced scattering coefficient and *conc*. is the concentration of intralipid. These parameters were also summarized in Supplementary Table 1.

The illumination waist measured in water at NA = 0.039 was used as initial FWHM of incident light in Monte Carlo simulations. The simulated results were consistent with the experimental observations (Supplementary Figs 8 and 9). Generally, the length over which the light sheet transmits by less than 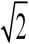times the initial waist (*w*0) is regarded as the distance useful for imaging ^52^. Under this definition, the critical length was larger than 1000 μm for the 1319-nm excitation in 1.25%, 2.50% and 5.00% intralipid solutions, much larger than that of 658-nm and 785-nm cases. Since the scattering coefficient of intralipid can be conveniently adjusted by controlling its concentration, it is a widely used phantom to study photon-material interactions. To study the light sheet propagation in an uniform scattering medium with similar scattering characteristics as mouse brain, we performed simulations using the scattering coefficient of the 2.5% intralipid and anisotropy factor of the brain measured by Shi *et al* ^53^. We compared the simulation results with our experimental observations in mouse brain in Supplementary Fig. 10. The simulated light propagation in brain was consistent with experimental results of 658-nm, 785-nm and 1,319-nm excitation (Supplementary Fig. 10f-h). The critical distances for uniform illumination were ~ 210 μm, ~ 320 μm and ~ 1000 μm for excitations using 658 nm, 785 nm and 1319 nm light sheet in mouse brain, respectively (Supplementary Fig. 10f-h).

The LS excitation intensity along incident direction is another important parameter for imaging in scattering tissue, as it affects the transmission distance of excitation in the tissue and determines the illumination field. As the intralipid concentration increased, the intensity along propagation direction attenuated faster but the 1319-nm excitation decayed the slowest compared to the 658-nm and 785-nm excitations (Supplementary Fig. 9j-l). Intensity attenuation was influenced by scattering, absorption and anisotropy of tissue. As the brain had larger anisotropy than intralipid, light sheet transmitted longer in the brain (Supplementary Fig. 10a,b,e). In order to study the illumination field of 658-nm, 785-nm and 1319-nm LS in mouse brain, we performed LSM imaging of a brain tissue at a depth of Z ~ 200 μm along the LS incident direction *X* for up to 1 cm (data in Fig. 2a,b). Though photons in the LS could propagate for as far as ~ 6000 μm to excite fluorescence in large-diameter vasculatures, only in the initial limited distance that small blood vessels (FWHM < 10 μm) could be observed (Fig. 2a). These limited distances were ~1380 μm, ~1676 μm and ~3900 μm for 658-nm, 785-nm and 1319-nm excitation respectively as longer excitation attenuated slower in mouse brain.

For high-quality optical sectioning in LSM, both uniform light sheet waist and available illumination field need to be ensured across the field of view.

## Acknowledgments

This study was supported by the National Institutes of Health NIH DP1-NS-105737.

## Author contributions

H.D., F.W., H.W. and J.Y. conceived and designed the experiments. H.D. and F.W. designed the optical system. F.W. set up the optical system. F.W., H.W., J.Y., H.M. and Z.M. performed the experiments. M.Z. developed the PbS/CdS CSQD dyes. Q.S. did the Monte Carlo simulations. F.W., H.W., J.Y., H.M., Z.M., Q.S., L.Q., Y.Z., Y.T., G.H., W.J.L., Y.L., L.L. and H.D. analyzed the data. F.W., W.H. and H.D. wrote the manuscript. All authors contributed to the general discussion and revision of the manuscript.

## Materials & Correspondence

Correspondence and requests for materials should be addressed to H.D.

